# Hornbills as drivers of plant range shifts under climate change

**DOI:** 10.1101/2025.10.06.680629

**Authors:** Vishesh L. Diengdoh

## Abstract

Range shifts of plants under climate change depend on frugivore-mediated seed dispersal. However, the role of frugivores in facilitating these shifts remains uncertain. This study assessed how plant-hornbill functional connectivity varied across latitude and elevation in South Asia by integrating species distribution models under present and future climate change scenarios with connectivity analysis under conservative (single-generation) and optimistic (multi-generational) seed dispersal scenarios. The connectivity patterns were synthesised using generalised additive models and indicated that latitude explained significantly more variation in connectivity than elevation. The range shifts of plants were primarily constrained by dispersal limitation rather than by the availability of suitable future habitat. Under conservative dispersal, connectivity remained stagnant despite climate change whereas optimistic dispersal fundamentally reshaped connectivity patterns, creating new pathways that aligned with areas of habitat gain. These findings demonstrate that the capacity of tropical plants to track climate change is critically dependent on hornbills, highlighting the urgent need to conserve both frugivore populations and landscape connectivity to ensure ecosystem resilience.

## Introduction

Climate change is driving shifts in species’ geographic ranges (Chen et al. 2011; Mason et al. 2015; Girish and Srinivasan 2022), however, the extent to which species can track these shifts depends not only on abiotic suitability but also on ecological interactions (Årevall et al. 2018; Lawlor et al. 2024). For plants, which are sessile, seed dispersal is a critical process enabling successful colonisation of new habitats (Rogers et al. 2021; Beckman and Sullivan 2023).

In tropical ecosystems, where 60–75% of plant species rely on animal-mediated dispersal (Rogers et al. 2021), range shifts in plant distributions are dependent on the persistence and movement of frugivorous animals. The loss of dispersal partners through defaunation will have cascading consequences on ecosystem functions and constrain plant responses to climate change (Fricke et al. 2022; Mendes et al. 2024; Svenning et al. 2024).

Hornbills (Bucerotidae) are among the largest frugivores in the tropical forests of Asia and Africa (Tobias et al. 2022), capable of dispersing seeds over distances exceeding 11 km (Naniwadekar et al. 2019) and interact with more than 470 plant species in Asia (Liang et al. 2024). These traits make them critical seed dispersers (Pires et al. 2018; Nowak et al. 2022; Fell et al. 2023). However, 81% of Asian hornbills are threatened to different degrees (IUCN 2025) raising concern that their decline could severely restrict plant responses to climate change.

Species distribution models (SDMs; Elith and Leathwick 2009) have greatly advanced the ability to project range shifts under climate change but are generally based on determining abiotic climatic suitability (e.g., Pang et al. 2024; Dyderski et al. 2025). Omniscape, a resistance based movement modelling framework based on circuit theory, estimates potential movement across landscapes by accounting for habitat and dispersal thresholds (McRae et al. 2008; Landau et al. 2021). Although widely applied to animal movement (e.g., Schloss et al. 2022; Gelmi Candusso et al. 2025), its potential for assessing the biotic facilitation of range shifts remains largely unexplored. Overall, understanding the potential for frugivores for facilitating range shifts requires integrating species distribution models with seed dispersal mechanisms.

This study assessed how plant-hornbill functional landscape connectivity varies with latitude and elevation across South Asia by developing a framework that combines projections of species distributions for plants and hornbills under present and future climate change scenarios with connectivity analysis under a conservative (static single-generation) and an optimistic (multi-generational expansive) seed dispersal scenarios. The findings reveal that the potential for plant range shifts is highly sensitive to which dispersal scenario is realised. The findings indicate that conserving functional landscape connectivity is essential for enabling the long-distance seed dispersal processes that underpin climate-driven range shifts.

## Methods

### Study Species

Plant–hornbill species and interactions were determined from (Liang et al. 2024). Plant species were restricted to non-introduced species occurring in tropical Asia (Govaerts et al. 2021; POWO 2025), where they overlap with hornbill distributions and are likely to depend on hornbills for climate-driven range shifts.

Occurrence records for both plants and hornbills were downloaded from Global Biodiversity Information Facility (GBIF; GBIF.Org 2025a, b). Records were limited to those after 1981. Duplicate records were removed. Spatial autocorrelation was reduced by keeping 1 record per pixel (1 km²).

Pseudo-absences were generated using the target-group method (Phillips et al. 2009) i.e., the presence records of other species were used as pseudo-absences. These pseudo-absences were ≥ 1 km from each other and all occurrence/presence points and the number of pseudo-absences equaled the number of presences.

Species with ≥ 100 cleaned presence and pseudo-absence records were retained for analysis, as this was assumed to be adequate for robust model training and testing.

### Study area

For each species, the study area was defined as the known range plus a 100 km buffer to account for potential range shifts. Plant ranges were obtained from (Govaerts et al. 2021; POWO 2025) while hornbill ranges were from the IUCN range data (IUCN 2025).

### Species Distribution Models

Species distributions were modelled using CHELSA bioclimatic variables (Karger et al. 2017, 2018) and Shuttle Radar Topography Mission (SRTM) elevation (Reuter et al. 2007; Jarvis et al. 2008) as predictors. Bioclimatic variables from 1981–2010 and 2011–2040 (SSP3–7.0, selected as the best representation of recent observed trends) represented present conditions, while 2070–2100 (SSP3–7.0 only for computational feasibility, across five GCMs) represented future scenarios.

Multicollinearity among predictors was reduced by excluding highly correlated variables (≥ 0.7) using a stepwise variance inflation factor (VIF) procedure in the *usdm* R package (Naimi et al. 2014).

The data were split into 70% training and 30% testing sets. Cross-validation was conducted using the training data for algorithm optimisation and for determining the probability threshold for binarisation via the *caret* R package (Kuhn 2008). The algorithms used include: artificial neural network, gradient boosting machine, k-nearest neighbour, random forest, and support vector machine.

The trained and optimised algorithms were used for projecting species distributions under present conditions and future scenarios; these outputs were averaged (unweighted) to result in the ensemble SDMs.

The accuracy of the cross-validated models were assessed using the Area Under the Curve (AUC) and True Skill Statistic (TSS). Similarly, AUC and TSS were calculated for the (present condition) ensemble models using the testing data.

The thresholds for converting probabilistic (0.0–1.0) ensemble projections to binary (0/1) were obtained using the sensitivity–specificity method (Cantor et al. 1999; Manel et al. 2001; Liu et al. 2005), averaged across cross-validation runs.

### Dispersal distance

Hornbill dispersal distance was estimated from gut passage time (GPT), flight speed (FS), and body mass (BM) following the trait-based allometric equations by Sorensen et al. (2020):

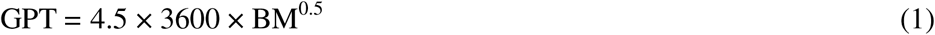

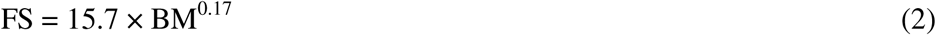

Seed dispersal distance (d) is the product of GPT, FS, BM, and a calibration constant (fc):

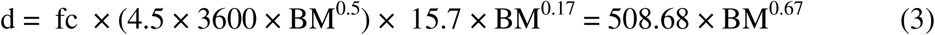

The value for BM was obtained from Tobias et al. (2022). Assuming that dispersal occurs continuously over time, cumulative dispersal distance (cd) over 75 years (2025–2100) was calculated using a power-law scaling (exponent 0.75) to reflect sublinear temporal accumulation:

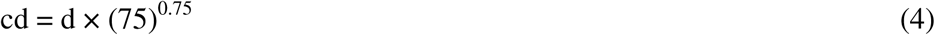

This framework resulted in two dispersal metrics: dispersal distance (d) which represented conservative static short-term single generation range shift potential (applied to both present and future connectivity models) and cumulative dispersal distance (cd) which represented optimistic multi-generational expansive range shift potential (applied only to future connectivity models; Supplementary Information (SI) Table 1)

**Table 1.**
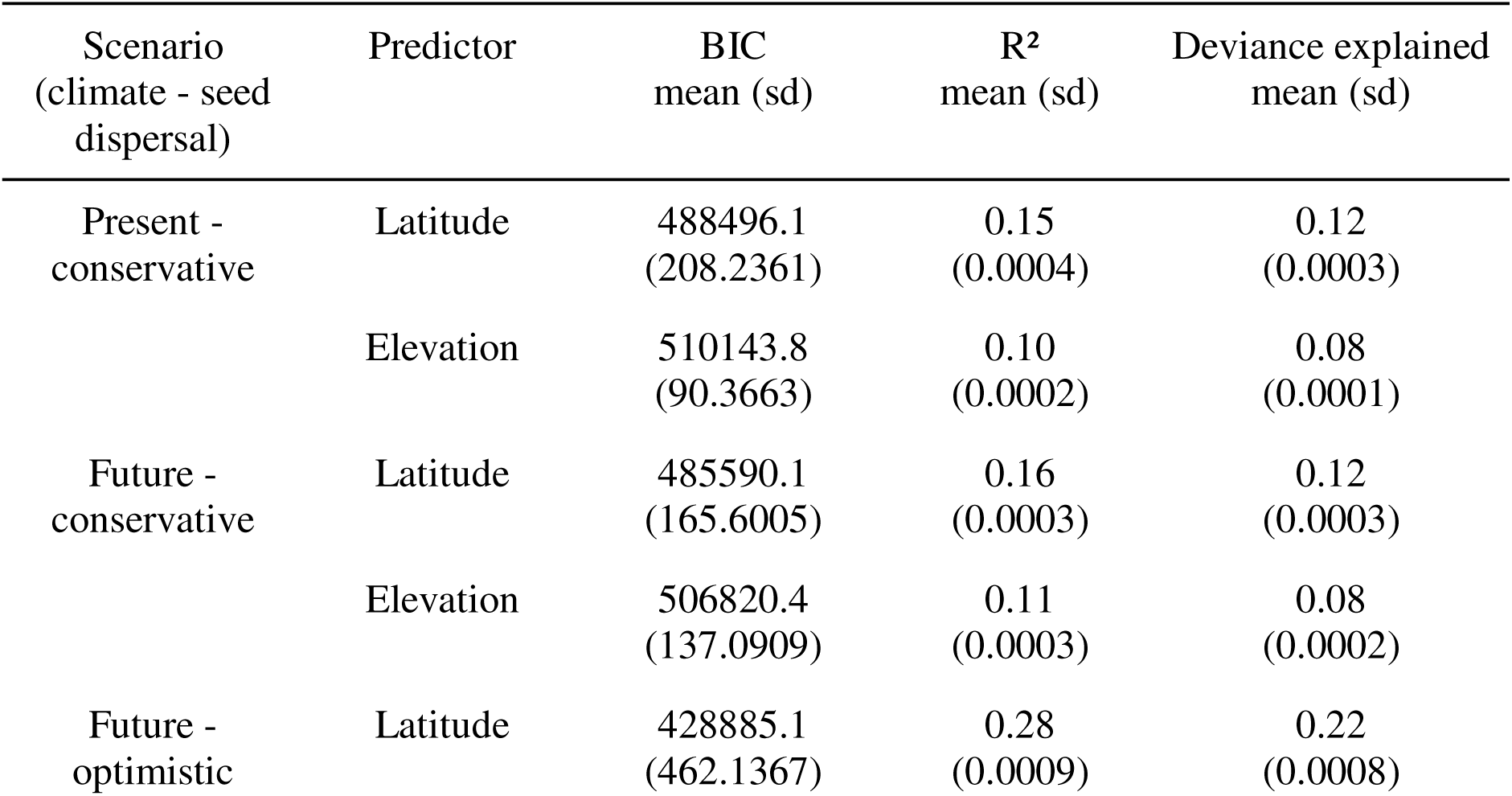
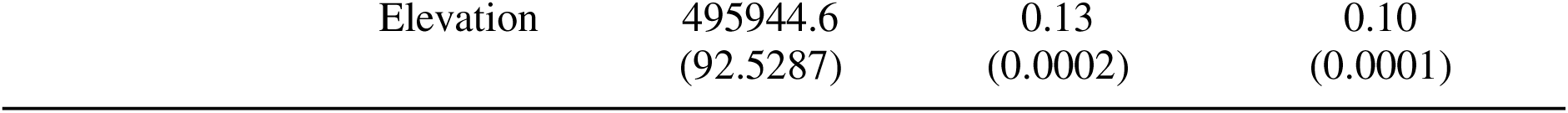
The bootstrapped Bayesian Information Criterion (BIC), adjusted R², and deviance explained mean and standard deviation (sd) values of the fitted Generalised Additive Models (GAMs) for different predictors and under climate and seed dispersal scenarios.

### Connectivity models

Landscape functional connectivity of each plant–hornbill interaction was estimated using the *Omniscape* Julia package (Landau et al. 2021). Omniscape required three inputs (source layer, resistance layer, radius) and produced two outputs (flow potential and normalised cumulative current).

The source layer was the binary SDM of each plant species. Under present conditions, only suitable habitat was used as the source. While under future scenarios stable, gained, and lost habitats were used. This approach assumes that species are able to disperse from habitat projected to become lost into stable or gained habitat before local extirpation occurs as well as from stable to gained habitat, thereby facilitating range shift.

The resistance layer was based on probabilistic SDM of hornbills where high probability equaled low resistance and vice versa. For present conditions, the present SDM was used, while for future scenarios the mean of present and future SDMs was used.

The radius defines the pixel-based moving window for identifying source–target connections. As Omniscape uses pixels rather than distance units, the radius was set to dispersal distance/1000 (each pixel is 1 km^2^; SI Table 1). The present connectivity models were parameterised using conservative dispersal distance, while future models were run under both conservative and optimistic dispersal distances.

The normalised cumulative current output was calculated as raw current divided by flow potential (null current when all landscape resistance = 1). Normalised cumulative current is a proxy for species movement where higher current indicates higher probability of movement.

The accuracy of the models were assessed by comparing differences in cumulative current between the presence and pseudo-absences points (using the testing data used to assess the accuracy of the species distribution models). If the landscape promoted connectivity, then the presence points should, on average, have higher values than pseudo-absences (Grafius et al. 2017; Rodrigues et al. 2022).

### Generalised Additive Models (GAMs)

Generalised Additive Models were implemented using the *mgcv* R package (Wood 2003, 2011, 2017) to assess the effect of latitude and elevation on the occurrence of connectivity areas (i.e., areas with cumulative current ≥ 1) under present and future climate change scenarios and conservative optimistic seed dispersal scenarios.

The response variable (connectivity occurrence) was modelled with a binomial distribution and logit link. The predictors (latitude or elevation) were fitted using penalised thin plate regression splines with shrinkage (“ts”). To account for non-independence among species, the model included a random effect for plant–hornbill interaction identity, implemented as a factor smooth (“re”).

Initial models included both latitude and elevation, but these predictors exhibited high concurvity. To avoid issues of collinearity and to enable a clear interpretation of each variable’s individual effect, separate models were fitted for latitude and elevation:

1. connectivity ∼ s(latitude, bs = ‘ts’, k= k) + s(interaction, bs = “re”)
2. connectivity ∼ s(elevation, bs = ‘ts’, k= k) + s(interaction, bs = “re”)

The model fit was assessed via inspection of residual plots, checks for concurvity, and comparison of Bayesian Information Criterion (BIC; stricter penalty for model complexity). The explanatory power was evaluated using the adjusted R^2^ and proportion of deviance explained.

All analyses were conducted in R v4.4.0 (R Core Team 2025). To facilitate transparency and reproducibility, the data and R code is available on the Open Science Framework: https://osf.io/m8g4f/?view_only=f869ae0bed6a4522a344ecaa68f28cb8.

## Results

Species distribution models were projected for 27 plant species and 16 hornbill species under present conditions of climate change. The differences in AUC and TSS metrics between the (averaged) cross-validated and ensemble models were minimal indicating good model fit (SI Table 2-3).

Functional landscape connectivity analysis was estimated for 60 plant–hornbill interactions under present conditions. Since 10 interactions did not meet accuracy threshold (mean cumulative current of presence points < pseudo-absence points) the remaining 50 interactions were retained for further analyses (SI Table 4).

The projected gains in habitat show limited overlap between hornbills and plants across latitude (Fig. 1b), with hornbills concentrated near the equator while plants gain habitat primarily north of the equator. However, projected stable and lost habitats show high overlap across environmental gradients (Fig 1a,c,d,f). Overall, these patterns suggest a decoupling in future expansion opportunities between hornbills and plants but a shared vulnerability in terms of habitat loss.

**Fig. 1.**
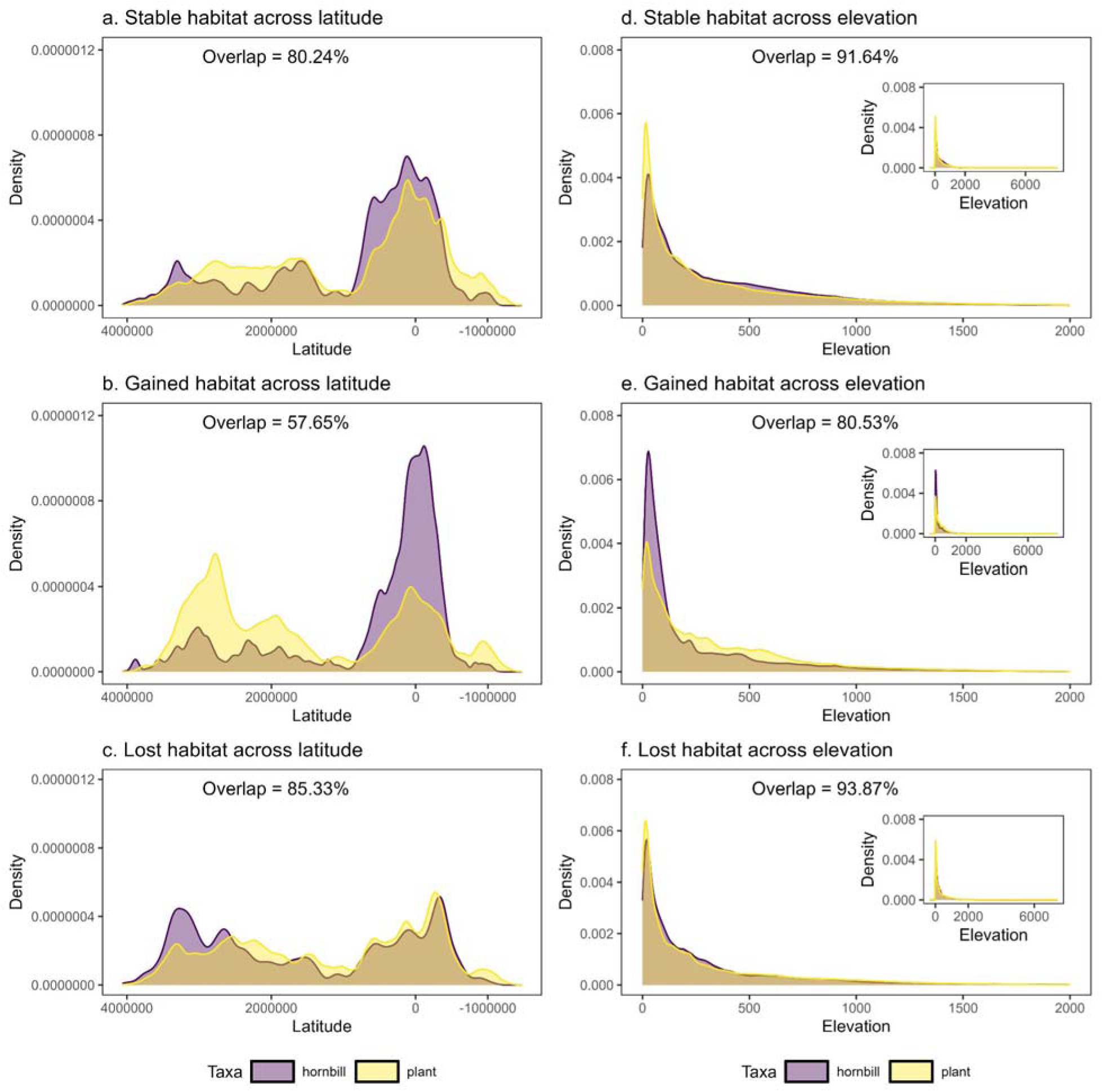
The density and overlap of the projected stable (a, d), gained (b, e), and lost (c, f) habitat between hornbills and plants across latitude and elevation. The overlap percentage was calculated using the *overlapping* R package (Pastore 2018).

The connectivity models projected the highest levels of cumulative current under the future climate scenario combined with optimistic seed dispersal (mean ± SD = 0.509 ± 0.497).

Connectivity values for the conservative dispersal scenario were low and showed minimal change between the present (0.323 ± 0.417) and future (0.334 ± 0.419) climate scenarios. This indicates that dispersal capacity, not climate-driven habitat shifts alone, is the primary determinant of range shift potential.

The results from the generalised additive models revealed that models with latitude as a predictor exhibited lower BIC scores, higher adjusted R², explained a greater proportion of deviance (Table 1) and had narrower confidence intervals than the elevation models (Fig 2b, 3b).

**Fig. 2.**
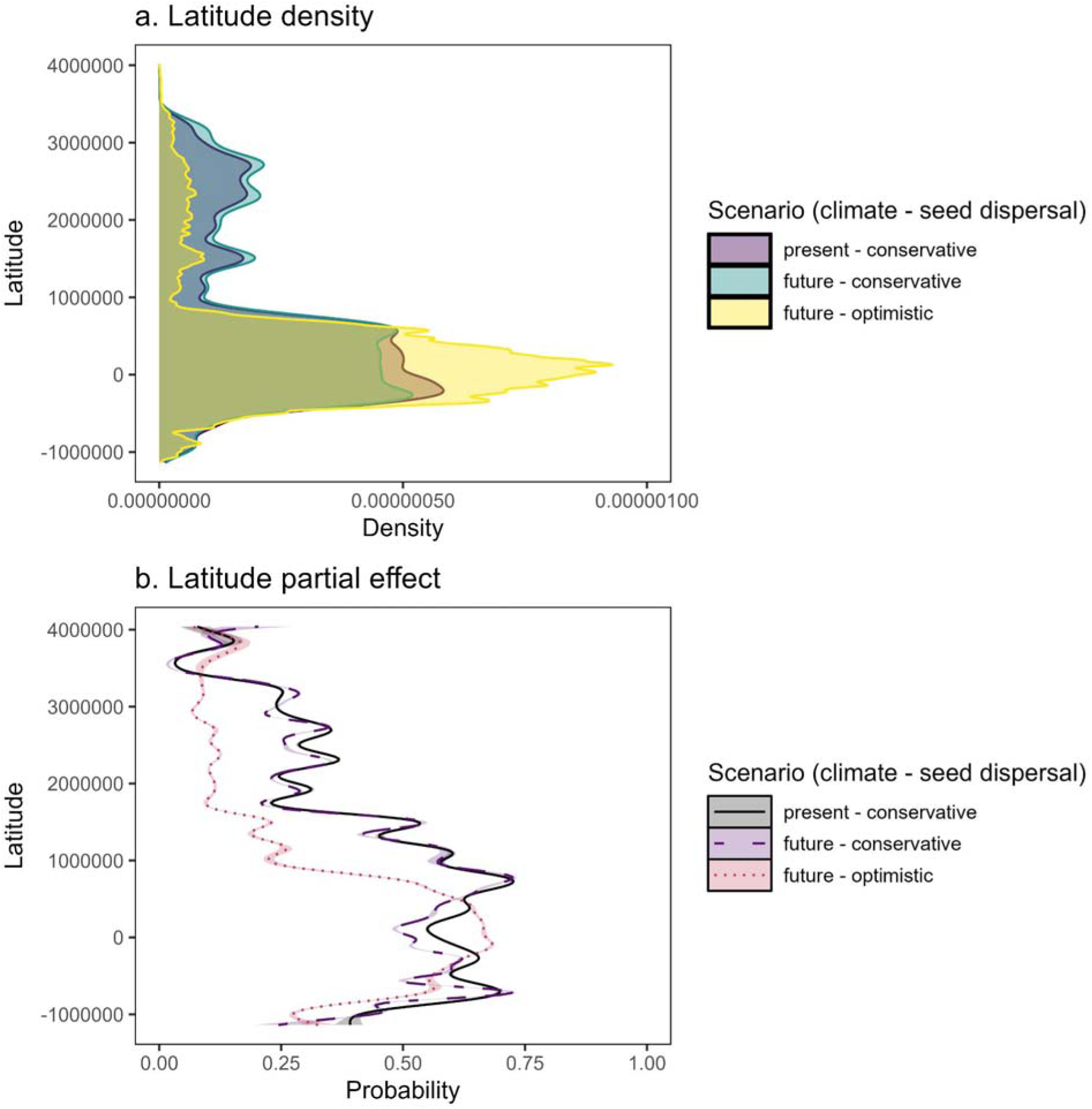
The (a) density of connectivity areas across latitude (projected coordinate system) and (b) partial effects of latitude (projected coordinate system) on occurrence of connectivity areas under present and future climate change and conservative and optimistic seed dispersal scenarios. In (b) the solid lines show bootstrapped fitted smooths from generalised additive models, with shaded ribbons representing 95% confidence intervals.

Although the latitude models were statistically better, the results for both predictors are presented because of their ecological importance in shaping species distributions.

Under present and future-conservative dispersal scenarios, both the density and probability of connectivity exhibited a pronounced non-linear pattern, peaking at mid-latitudes (∼0 to 2 million m – projected coordinate system; Fig. 2a,b). This suggests that connectivity was strongly constrained within a specific latitudinal band under conservative dispersal scenarios. The future-optimistic dispersal scenario, however, fundamentally reorganises this spatial pattern. The density plot shows a secondary peak emerging at high latitudes (∼3 to 4 million m – projected coordinate system; Fig. 2a), and this northward shift is captured with high confidence but low probability by the partial effect plot (Fig. 2b). The close alignment between the raw density data and the modeled partial effect underscores the robustness of these findings. This latitudinal shift in connectivity under optimistic dispersal aligns geographically with the predominant areas of habitat gain for plant species (Fig. 1b), indicating that enhanced dispersal capacity enables the realisation of range-shift potential into newly available climatic niches.

The relationship between connectivity and elevation was relatively weaker and less structured than for latitude (Fig. 3). The partial effect plot for elevation confirms its limited explanatory power, showing low and relatively invariant probability across the gradient (Fig. 3b). However, model predictions below 2000 meters are reliable because this range is well-supported by the data. Within this reliable range, the probability of connectivity was consistently highest at lower elevations across all scenarios, indicating that lowland areas are the core zones for connectivity. The persistence of this pattern across dispersal scenarios suggests that the concentration of connectivity at lower elevations is a fundamental feature of the system, not a consequence of dispersal limitation.

**Fig. 3.**
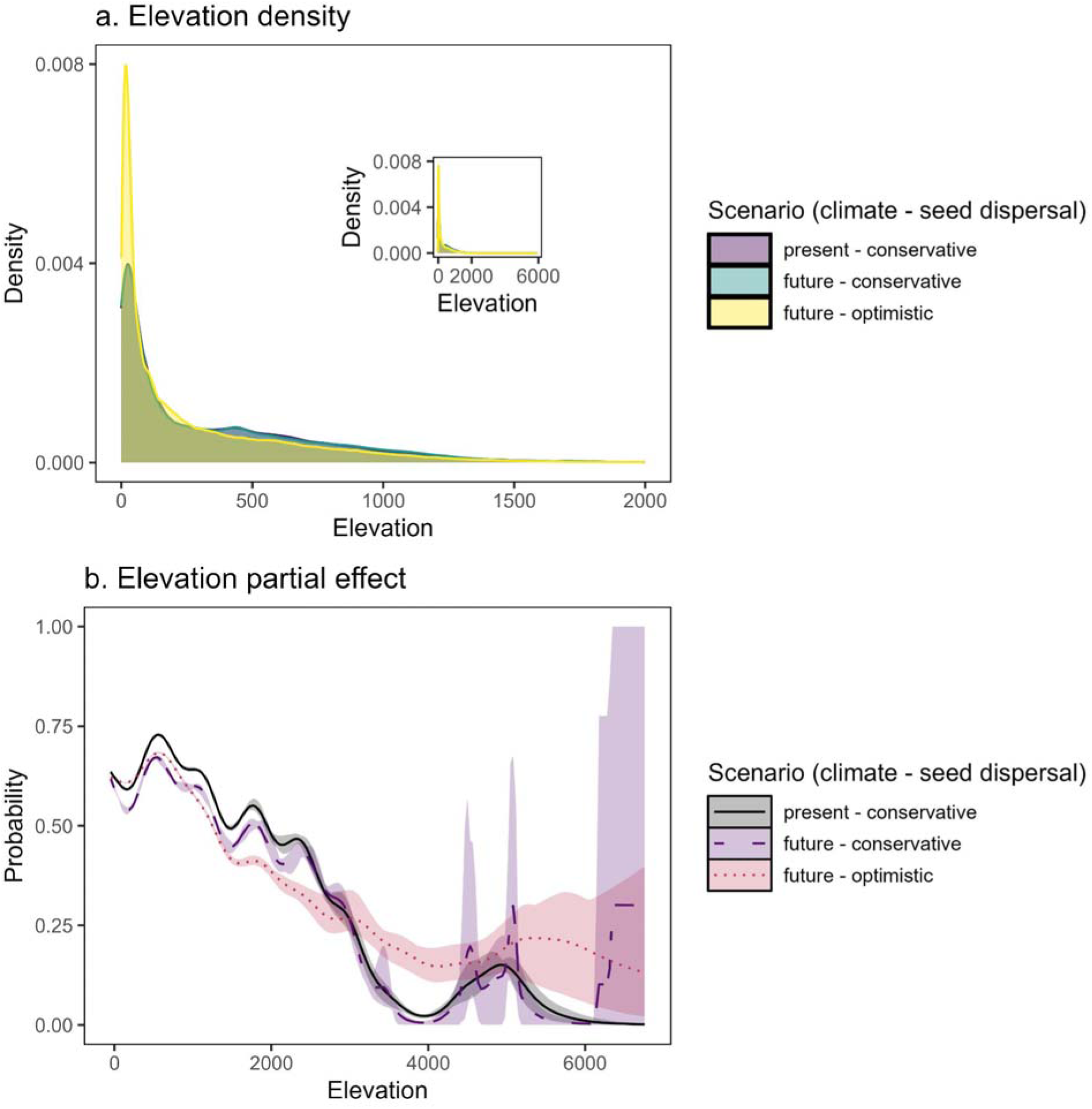
The (a) density of connectivity areas across elevation and (b) partial effects of elevation (m above sea level) on occurrence of connectivity areas under present and future climate change and conservative and optimistic seed dispersal scenarios. In (b) the solid lines show bootstrapped fitted smooths from generalised additive models, with shaded ribbons representing 95% confidence intervals.

## Discussion

By integrating species distribution models with a trait-based dispersal connectivity analysis, this study (i) indicates latitude is a statistically better than elevation in explaining connectivity and (ii) projects that the potential for hornbills to facilitate plant range shifts under climate change is constrained more by dispersal limitation than by the availability of future suitable habitat. These findings highlight that broad-scale latitudinal processes and biotic interactions will play a decisive role in shaping plant responses to climate change.

The stronger statistical performance of latitude over elevation suggests that continental-scale patterns of connectivity and in turn range shifts are primarily governed by latitudinal gradients. This contrasts with many studies observing elevational range shifts, particularly in mountains (Mamantov et al. 2021; Girish and Srinivasan 2022) and tropical countries (Colwell and Feeley 2025). These apparently conflicting results can be reconciled through a scale-dependent framework: at local and mountain-system scales, elevation is the dominant driver, while at biogeographic scales, latitude emerges as the stronger gradient. This highlights the importance of explicitly considering scale when interpreting climate-driven distributional changes.

Under the optimistic dispersal scenario, connectivity reorganises into a bimodal distribution, with pronounced peaks at both equatorial and high latitudes corresponding to areas of projected habitat gain. This unrealised potential highlights the critical role of long-distance dispersal in enabling species to track shifting climates. This high-latitude connectivity is consistent with poleward range shifts commonly documented in temperate ecosystems (Svenning 2015; Rubenstein et al. 2023) but rarely observed in the tropics, where data are scarce. In South and Southeast Asia, such northward shifts are likely facilitated by the contiguous landmass that provides a pathway for gradual dispersal. In contrast, the insular regions of the Indo-Malayan archipelago (e.g., Borneo, Java) lack such opportunities because of oceanic barriers. This mainland–island dichotomy emphasises that conservation strategies must be region-specific: maintaining transnational latitudinal corridors across South Asia, particularly along the large landmass of the Indo-Burma biodiversity hotspot is vital for facilitating range shifts, while safeguarding island forests is essential for preserving persistent strongholds.

The stagnation of connectivity under conservative dispersal scenarios highlights a severe dispersal limitation bottleneck, indicating that long distance multigenerational seed dispersal is essential for keeping pace with climate change (Schloss et al. 2012; Nowak et al. 2022).

Across all scenarios, connectivity consistently concentrates in lowland forests. This persistence indicates that lowlands form the stable ecological backbone of hornbill–plant interactions, rather than an artefact of model assumptions. Importantly, this finding aligns with current observations of limited upslope shifts in tropical mountain systems (Chen et al. 2025; Farfan-Rios et al. 2025) suggesting that the potential for elevational escape is highly constrained and will continue to be so. Conservation of lowland corridors is therefore critical, not only to sustain present ecological networks but also to secure future pathways for range shifts.

The study has limitations which also provide a clear direction for future research. The spatial scale of 1 km² masks fine-scale topographic heterogeneity, particularly in the complex mountain systems. As a result, localised elevational shifts are not captured, underscoring the need for complementary high-resolution studies at local scales. The dispersal distances for the connectivity models are derived from allometric equations rather than empirical movement data. While GPS tracking of hornbills are available for some species (see Naniwadekar et al. 2019) there is a pressing need to have tracking data for more species and across climatic seasons and landscapes; these would dramatically improve connectivity forecasts and strengthen conservation planning.

In conclusion, this study demonstrates that range shifts for plants in South Asia are dependent on hornbills. Although potential pathways for future range shifts exist, they are locked by dispersal limitations. The optimistic dispersal scenario where connectivity expands northward and increases at the equator as well as the persistence of connectivity at low elevations across scenarios can be realised through urgent and targeted conservation action – conservation of transnational latitudinal corridors and lowland forests. Given that many Asian hornbill species are already threatened (IUCN 2025) and that habitat loss and fragmentation represent some of the most severe biodiversity threats in the region (Namkhan et al. 2021; Struebig et al. 2025), conservation strategies must also explicitly address both hornbill populations and their habitats. Safeguarding future plant diversity is directly linked to protecting viable hornbill populations and connectivity corridors today.

## Author Contributions

VLD: Conceptualisation, Methodology, Analysis, Visualisation, Writing (original draft), Writing (review and editing).

## Data Availability

The results: species distribution models and connectivity models are accessible via Open Science Framework: https://osf.io/m8g4f/?view_only=f869ae0bed6a4522a344ecaa68f28cb8

## Code Availability

All R code used for data analysis is provided and hosted on Open Science Framework: https://osf.io/m8g4f/?view_only=f869ae0bed6a4522a344ecaa68f28cb8

