## Supplementary Information for "Hornbills as drivers of plant range shifts under climate change"

**Supplementary Information (SI)**

**SI Table 1** The dispersal and cumulative dispersal distances and radii of hornbill species.

| SI | Species | Mass (kg) | Dispersal  (m) | Radius* (dispersal) | Cumulative dispersal  (m) | Radius (cumulative dispersal) |
| --- | --- | --- | --- | --- | --- | --- |
| 1 | *Aceros nipalensis* | 2.385 | 910.6717814 | 1 | 23209 | 23 |
| 2 | *Anorrhinus austeni* | 0.933 | 485.5851132 | 1 | 12375 | 12 |
| 3 | *Anorrhinus galeritus* | 1.172 | 565.7520841 | 1 | 14419 | 14 |
| 4 | *Anorrhinus tickelli* | 0.7926 | 435.322273 | 1 | 11094 | 11 |
| 5 | *Anthracoceros albirostris* | 0.8929 | 471.5009303 | 1 | 12017 | 12 |
| 6 | *Anthracoceros coronatus* | 0.8067 | 440.4957684 | 1 | 11226 | 11 |
| 7 | *Anthracoceros malayanus* | 1.05 | 525.5832221 | 1 | 13395 | 13 |
| 8 | *Berenicornis comatus* | 1.47 | 658.4871083 | 1 | 16782 | 17 |
| 9 | *Buceros bicornis* | 2.7908 | 1011.77257 | 1 | 25786 | 26 |
| 10 | *Buceros rhinoceros* | 2.3716 | 907.2404985 | 1 | 23122 | 23 |
| 11 | *Ocyceros birostris* | 0.375 | 263.6608282 | 1 | 6720 | 7 |
| 12 | *Ocyceros gingalensis* | 0.238 | 194.4246043 | 1 | 4955 | 5 |
| 13 | *Ocyceros griseus* | 0.292 | 222.9724332 | 1 | 5683 | 6 |
| 14 | *Rhabdotorrhinus corrugatus* | 1.59 | 694.0339829 | 1 | 17688 | 18 |
| 15 | *Rhinoplax vigil* | 2.8876 | 1035.152832 | 1 | 26382 | 26 |
| 16 | *Rhyticeros undulatus* | 2.2146 | 866.5477734 | 1 | 22085 | 22 |

***** Radius was set to 1 the minimum possible radius size for species where dispersal is less than 1 km.

**SI Table 2** The AUC and TSS metrics of the cross validated (CV) and ensemble models of plant species.

| SI | Species | CV  AUC | CV  TSS | Ensemble AUC | Ensemble TSS |
| --- | --- | --- | --- | --- | --- |
| 1 | *Adenia macrophylla* | 0.84 | 0.60 | 0.75 | 0.23 |
| 2 | *Artocarpus elasticus* | 0.75 | 0.45 | 0.73 | 0.19 |
| 3 | *Artocarpus hirsutus* | 0.89 | 0.75 | 0.96 | 0.88 |
| 4 | *Dacryodes rostrata* | 0.81 | 0.55 | 0.83 | 0.45 |
| 5 | *Diospyros montana* | 0.76 | 0.47 | 0.82 | 0.48 |
| 6 | *Euonymus indicus* | 0.75 | 0.44 | 0.84 | 0.48 |
| 7 | *Ficus deltoidea* | 0.74 | 0.37 | 0.72 | 0.31 |
| 8 | *Ficus mollis* | 0.63 | 0.29 | 0.58 | 0.13 |
| 9 | *Ficus sumatrana* | 0.66 | 0.33 | 0.60 | 0.18 |
| 10 | *Ficus sundaica* | 0.78 | 0.49 | 0.77 | 0.39 |
| 11 | *Ficus villosa* | 0.74 | 0.41 | 0.81 | 0.48 |
| 12 | *Gironniera nervosa* | 0.78 | 0.50 | 0.77 | 0.44 |
| 13 | *Gymnacranthera farquhariana* | 0.75 | 0.49 | 0.66 | 0.22 |
| 14 | *Knema latericia* | 0.82 | 0.58 | 0.82 | 0.61 |
| 15 | *Knema laurina* | 0.55 | 0.19 | 0.82 | 0.53 |
| 16 | *Macaranga gigantea* | 0.62 | 0.29 | 0.75 | 0.43 |
| 17 | *Macaranga peltata* | 0.90 | 0.71 | 0.94 | 0.72 |
| 18 | *Madhuca longifolia* | 0.69 | 0.32 | 0.82 | 0.48 |
| 19 | *Microcos tomentosa* | 0.73 | 0.46 | 0.68 | 0.32 |
| 20 | *Polyalthia cauliflora* | 0.77 | 0.44 | 0.82 | 0.51 |
| 21 | *Prunus arborea* | 0.68 | 0.36 | 0.81 | 0.59 |
| 22 | *Schleichera oleosa* | 0.73 | 0.38 | 0.66 | 0.16 |
| 23 | *Sterculia guttata* | 0.78 | 0.51 | 0.90 | 0.63 |
| 24 | *Tetrapilus dioicus* | 0.92 | 0.76 | 0.89 | 0.63 |
| 25 | *Trichosanthes tricuspidata* | 0.60 | 0.25 | 0.63 | 0.26 |
| 26 | *Uvaria littoralis* | 0.77 | 0.48 | 0.68 | 0.32 |
| 27 | *Vitex altissima* | 0.76 | 0.54 | 0.65 | 0.20 |

**SI Table 3** The AUC and TSS metrics of the cross validated (CV) and ensemble models of hornbill species.

| SI | Species | CV AUC | CV TSS | Ensemble AUC | Ensemble TSS |
| --- | --- | --- | --- | --- | --- |
| 1 | *Aceros nipalensis* | 0.83 | 0.56 | 0.90 | 0.64 |
| 2 | *Anorrhinus austeni* | 0.88 | 0.67 | 0.92 | 0.67 |
| 3 | *Anorrhinus galeritus* | 0.78 | 0.45 | 0.83 | 0.49 |
| 4 | *Anorrhinus tickelli* | 0.88 | 0.66 | 0.93 | 0.76 |
| 5 | *Anthracoceros albirostris* | 0.81 | 0.50 | 0.86 | 0.56 |
| 6 | *Anthracoceros coronatus* | 0.84 | 0.55 | 0.88 | 0.58 |
| 7 | *Anthracoceros malayanus* | 0.78 | 0.44 | 0.82 | 0.48 |
| 8 | *Berenicornis comatus* | 0.82 | 0.53 | 0.82 | 0.51 |
| 9 | *Buceros bicornis* | 0.84 | 0.53 | 0.88 | 0.59 |
| 10 | *Buceros rhinoceros* | 0.80 | 0.48 | 0.84 | 0.53 |
| 11 | *Ocyceros birostris* | 0.75 | 0.36 | 0.79 | 0.42 |
| 12 | *Ocyceros gingalensis* | 0.74 | 0.37 | 0.72 | 0.29 |
| 13 | *Ocyceros griseus* | 0.92 | 0.71 | 0.92 | 0.72 |
| 14 | *Rhabdotorrhinus corrugatus* | 0.85 | 0.57 | 0.87 | 0.51 |
| 15 | *Rhinoplax vigil* | 0.77 | 0.48 | 0.82 | 0.54 |
| 16 | *Rhyticeros undulatus* | 0.82 | 0.52 | 0.86 | 0.56 |

**SI Table 4** The mean cumulative current of presence and pseudo-absence points indicating the accuracy of the connectivity models.

| SI | Plant | Hornbill | Presence  mean cumulative current | Pseudo-absence mean cumulative current | Accuracy* |
| --- | --- | --- | --- | --- | --- |
| 1 | *Adenia macrophylla* | *Anorrhinus galeritus* | 0.4551 | 0.2789 | 1 |
| 2 | *Artocarpus elasticus* | *Buceros bicornis* | 0.1411 | 0.0766 | 1 |
| 3 | *Artocarpus elasticus* | *Rhyticeros undulatus* | 0.2774 | 0.1465 | 1 |
| 4 | *Artocarpus hirsutus* | *Ocyceros griseus* | 0.5595 | 0.1535 | 1 |
| 5 | *Dacryodes rostrata* | *Anorrhinus galeritus* | 0.6414 | 0.4471 | 1 |
| 6 | *Dacryodes rostrata* | *Anthracoceros albirostris* | 0.2804 | 0.1646 | 1 |
| 7 | *Dacryodes rostrata* | *Anthracoceros malayanus* | 0.7763 | 0.4799 | 1 |
| 8 | *Dacryodes rostrata* | *Berenicornis comatus* | 0.5327 | 0.3397 | 1 |
| 9 | *Dacryodes rostrata* | *Buceros rhinoceros* | 0.6581 | 0.4385 | 1 |
| 10 | *Dacryodes rostrata* | *Rhabdotorrhinus corrugatus* | 0.7457 | 0.4658 | 1 |
| 11 | *Dacryodes rostrata* | *Rhinoplax vigil* | 0.5492 | 0.4574 | 1 |
| 12 | *Dacryodes rostrata* | *Rhyticeros undulatus* | 0.3846 | 0.1260 | 1 |
| 13 | *Diospyros montana* | *Buceros bicornis* | 0.3222 | 0.3914 | 0 |
| 14 | *Diospyros montana* | *Ocyceros birostris* | 0.3298 | 0.3391 | 0 |
| 15 | *Euonymus indicus* | *Ocyceros birostris* | 0.0351 | 0.0466 | 0 |
| 16 | *Ficus deltoidea* | *Buceros bicornis* | 0.0689 | 0.0284 | 1 |
| 17 | *Ficus deltoidea* | *Rhyticeros undulatus* | 0.2818 | 0.0920 | 1 |
| 18 | *Ficus mollis* | *Ocyceros gingalensis* | 0.4439 | 0.3535 | 1 |
| 19 | *Ficus sumatrana* | *Anthracoceros malayanus* | 0.2666 | 0.3131 | 0 |
| 20 | *Ficus sumatrana* | *Buceros rhinoceros* | 0.2390 | 0.3059 | 0 |
| 21 | *Ficus sundaica* | *Anorrhinus galeritus* | 0.3765 | 0.3486 | 1 |
| 22 | *Ficus sundaica* | *Anthracoceros malayanus* | 0.5173 | 0.3955 | 1 |
| 23 | *Ficus sundaica* | *Berenicornis comatus* | 0.2985 | 0.2881 | 1 |
| 24 | *Ficus sundaica* | *Buceros bicornis* | 0.1264 | 0.0849 | 1 |
| 25 | *Ficus sundaica* | *Buceros rhinoceros* | 0.3854 | 0.3529 | 1 |
| 26 | *Ficus sundaica* | *Rhabdotorrhinus corrugatus* | 0.6421 | 0.4293 | 1 |
| 27 | *Ficus sundaica* | *Rhinoplax vigil* | 0.2746 | 0.3542 | 0 |
| 28 | *Ficus sundaica* | *Rhyticeros undulatus* | 0.2013 | 0.1218 | 1 |
| 29 | *Ficus villosa* | *Anorrhinus austeni* | 0.2304 | 0.0330 | 1 |
| 30 | *Ficus villosa* | *Anorrhinus galeritus* | 0.7493 | 0.5296 | 1 |
| 31 | *Ficus villosa* | *Anthracoceros malayanus* | 0.7021 | 0.5594 | 1 |
| 32 | *Ficus villosa* | *Buceros rhinoceros* | 0.7244 | 0.5644 | 1 |
| 33 | *Gironniera nervosa* | *Anorrhinus austeni* | 0.0000 | 0.0000 | 0 |
| 34 | *Gironniera nervosa* | *Buceros bicornis* | 0.0625 | 0.0465 | 1 |
| 35 | *Gymnacranthera farquhariana* | *Rhyticeros undulatus* | 0.2593 | 0.1735 | 1 |
| 36 | *Knema latericia* | *Anorrhinus galeritus* | 0.5467 | 0.4248 | 1 |
| 37 | *Knema latericia* | *Rhyticeros undulatus* | 0.3560 | 0.1465 | 1 |
| 38 | *Knema laurina* | *Aceros nipalensis* | 0.0468 | 0.0070 | 1 |
| 39 | *Knema laurina* | *Anorrhinus austeni* | 0.0370 | 0.0000 | 1 |
| 40 | *Knema laurina* | *Anorrhinus tickelli* | 0.0719 | 0.0000 | 1 |
| 41 | *Knema laurina* | *Anthracoceros albirostris* | 0.2036 | 0.1397 | 1 |
| 42 | *Knema laurina* | *Buceros bicornis* | 0.1755 | 0.0617 | 1 |
| 43 | *Knema laurina* | *Rhyticeros undulatus* | 0.2915 | 0.1334 | 1 |
| 44 | *Macaranga gigantea* | *Buceros bicornis* | 0.1495 | 0.0734 | 1 |
| 45 | *Macaranga gigantea* | *Rhyticeros undulatus* | 0.2317 | 0.1346 | 1 |
| 46 | *Macaranga peltata* | *Ocyceros griseus* | 0.6250 | 0.2547 | 1 |
| 47 | *Madhuca longifolia* | *Anthracoceros coronatus* | 0.2328 | 0.5302 | 0 |
| 48 | *Microcos tomentosa* | *Rhabdotorrhinus corrugatus* | 0.2925 | 0.1821 | 1 |
| 49 | *Polyalthia cauliflora* | *Anorrhinus galeritus* | 0.5036 | 0.3847 | 1 |
| 50 | *Prunus arborea* | *Rhyticeros undulatus* | 0.3983 | 0.1940 | 1 |
| 51 | *Schleichera oleosa* | *Ocyceros birostris* | 0.3784 | 0.3373 | 1 |
| 52 | *Schleichera oleosa* | *Ocyceros gingalensis* | 0.0438 | 0.0385 | 1 |
| 53 | *Sterculia guttata* | *Ocyceros griseus* | 0.7305 | 0.3338 | 1 |
| 54 | *Tetrapilus dioicus* | *Buceros bicornis* | 0.1654 | 0.0999 | 1 |
| 55 | *Tetrapilus dioicus* | *Ocyceros griseus* | 0.5776 | 0.1813 | 1 |
| 56 | *Trichosanthes tricuspidata* | *Anorrhinus tickelli* | 0.8676 | 0.6077 | 1 |
| 57 | *Trichosanthes tricuspidata* | *Anthracoceros albirostris* | 0.4031 | 0.4327 | 0 |
| 58 | *Uvaria littoralis* | *Rhyticeros undulatus* | 0.4914 | 0.3481 | 1 |
| 59 | *Vitex altissima* | *Buceros bicornis* | 0.1110 | 0.1411 | 0 |
| 60 | *Vitex altissima* | *Ocyceros birostris* | 0.1240 | 0.0956 | 1 |

* Accuracy = 1 indicates interaction was retained for further analyses.
